# Comparative genomics identifies putative signatures of sociality in spiders

**DOI:** 10.1101/797241

**Authors:** Chao Tong, Gabriella M. Najm, Noa Pinter-Wollman, Jonathan N. Pruitt, Timothy A. Linksvayer

## Abstract

Comparative genomics has begun to elucidate the genomic basis of social life in insects but insight into the genomic basis of spider sociality has lagged behind. To begin to characterize genomic signatures associated with the evolution of social life in spiders, we performed one of the first spider comparative genomics studies including five solitary species and two social species, representing two independent origins of sociality in the genus *Stegodyphus*. We found that the two social spider species had a large expansion of gene families associated with transport and metabolic processes and an elevated genome-wide rate of molecular evolution compared with the five solitary spider species. Genes that were rapidly evolving in the two social species relative to the five solitary species were enriched for transport, behavior, and immune functions, while genes that were rapidly evolving in the solitary species were enriched for energy metabolism processes. Most rapidly evolving genes in the social species *S. dumicola* were broadly expressed across four tissues and enriched for transport functions, but 12 rapidly evolving genes showed brain-specific expression and were enriched for social behavioral processes. Altogether, our study identifies putative genomic signatures and potential candidate genes associated with spider sociality. These results indicate that future spider comparative genomic studies, including broader sampling and additional independent origins of sociality, can further clarify the genomic causes and consequences of social life.

## Introduction

A major goal of evolutionary biology is to elucidate the genomic underpinnings of phenotypic innovations across the tree of life. Over the past decade, comparative genomics has been used as a powerful tool to begin to elucidate the genetic basis of phenotypic innovations and adaptation in a wide range of organisms (Xu et al. 2017; Gou et al. 2014; Zhang et al. 2014; Chen et al. 2019; Exposito-Alonso et al. 2018; Gaither et al. 2018; Goodman et al. 2009; Wang et al. 2019; Wu et al. 2014). These studies have often identified changes in gene repertoire (e.g., expansions of certain gene families) or genomic signatures of molecular evolution at orthologous genes, by comparing the genomes of species with and without a phenotype of interest (Xu et al. 2017; Gou et al. 2014; Gaither et al. 2018; Goodman et al. 2009; Wang et al. 2019; Wu et al. 2014).

The evolution of group-living is a conspicuous phenotypic innovation found across diverse groups of animals, including many vertebrates, insects, and spiders (Rubenstein & Abbot 2017). A series of comparative genomic and transcriptomic studies in eusocial insects has begun to identify putative genomic signatures of the evolution of complex societies (Simola et al. 2013; Kapheim et al. 2015). These putative signatures include the expansion of certain gene families associated with functions such as chemical communication (Harrison et al. 2018; McKenzie et al. 2014, 2016) and signatures of elevated molecular evolution at certain genes (Roux et al. 2014; Privman et al. 2018; Jia et al. 2018; Kulmuni et al. 2013; Woodard et al. 2011). Studies in social insects and other social organisms have also commonly emphasized the importance of genes associated with metabolism (Woodard et al. 2011; Rittschof et al. 2014) and reproduction (Warner et al. 2019) as potentially making up a conserved social toolkit or groundplan (Toth & Robinson 2007; Amdam et al. 2006; Linksvayer & Wade 2005; Rittschof & Robinson 2016; Johnson & Linksvayer 2010).

While sociality has also evolved multiple times independently in spiders (Johannesen et al. 2007), the high complexity and large size of spider genomes has constrained the development of genomic resources for spiders (Schwager et al. 2017; Babb et al. 2017; Sanggaard et al. 2014; Liu et al. 2019; Garb et al. 2018), so that comparative genomic studies of spider sociality have lagged behind other groups such as social insects. Previous comparative genomic studies including spiders have compared newly sequenced spider genomes to available insect genomes (Sanggaard et al. 2014; Babb et al. 2017), but not across spiders or among other arachnids. These previous studies have mainly focused on identifying sets of venom and silk genes in spiders (Garb et al. 2018, 2019), but have also provided evidence for whole genome duplication and gene family expansion during arachnid evolution (Schwager et al. 2017; Babb et al. 2017; Sanggaard et al. 2014).

Building on previous comparative genomic work in social insects, we performed one of the first spider comparative genomic studies and focused in particular on identifying putative genomic signatures associated with the evolution of social life in spiders. Specifically, we used recently available genomes from seven spider species (Sanggaard et al. 2014; Liu et al. 2019; Babb et al. 2017; Schwager et al. 2017), including two social spiders with independent origins of sociality (Settepani et al. 2016; Johannesen et al. 2007) from the genus *Stegodyphus* (Liu et al. 2019; Sanggaard et al. 2014). We aimed to identify genome content and genome-wide patterns of molecular evolution that differ between social and solitary spiders. In addition, using new tissue-specific expression data we generated from the social spider *Stegodyphus dumicola*, we also characterized the expression pattern of genes identified as having elevated rates of molecular evolution in the two *Stegodyphus* social spider species.

## Methods

### Data retrieval and sequence analysis

We searched all available sequences, annotations, and predicted proteins of spider genomes and downloaded from public online databases (fig. 1A, supplementary table S1), including NCBI (https://www.ncbi.nlm.nih.gov/) and HGSC (https://www.hgsc.bcm.edu/). Specifically, we used the genomic data of the social spiders *Stegodyphus mimosarum* (v1.0, NCBI) (Sanggaard et al. 2014) and *S. dumicola* (Liu et al. 2019), and the solitary spiders *Parasteatoda tepidariorum* (v2.0, NCBI) (Schwager et al. 2017), *Acanthoscurria geniculata* (v1, NCBI) (Sanggaard et al. 2014), *Nephila clavipes* (v1.0, NCBI) (Babb et al. 2017), *Loxosceles reclusa* (v1.0, HGSC) and *Latrodectus hesperus* (v1.0, HGSC).

**FIG. 1.**
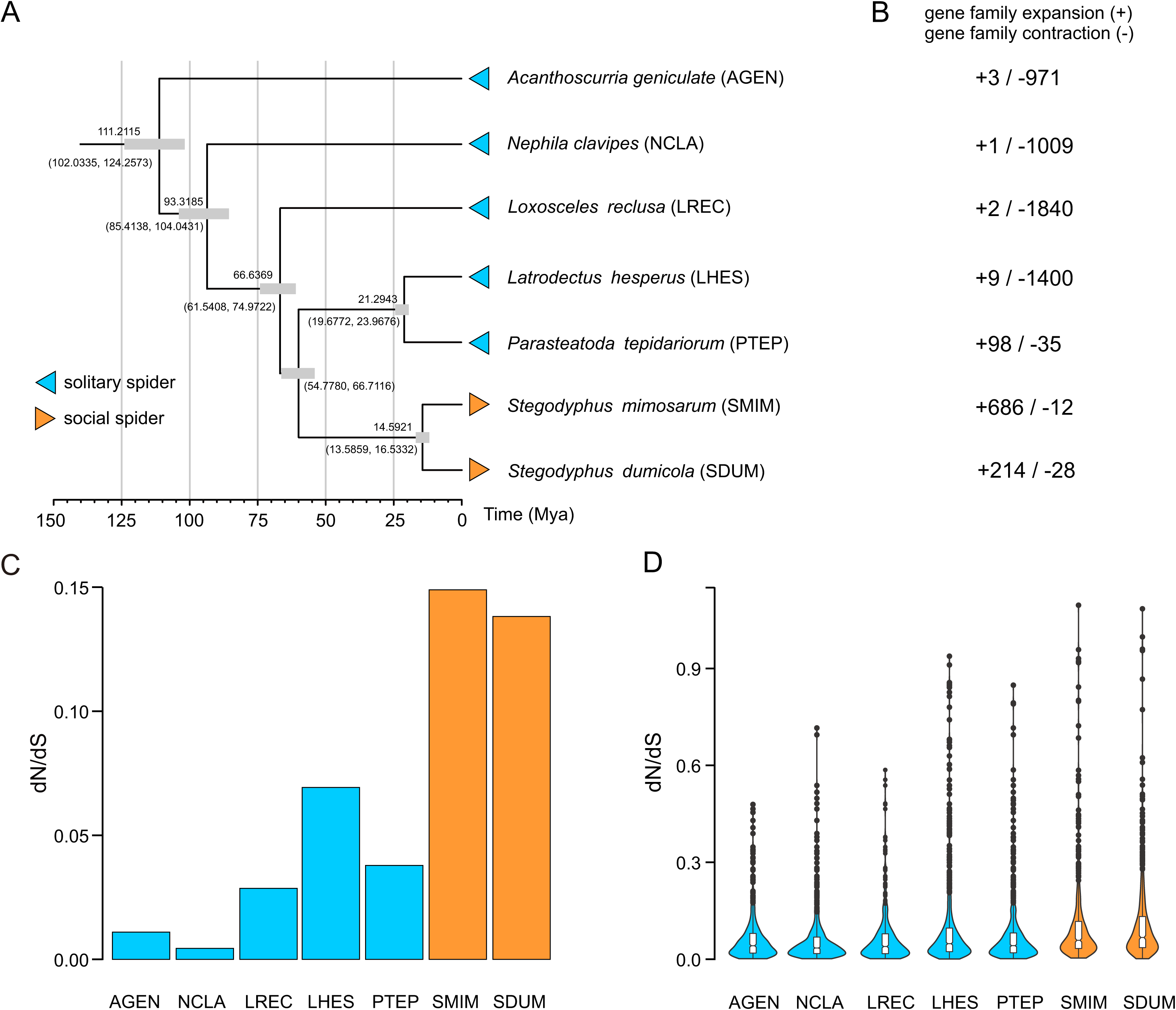
Genomic content and evolution in social spider relative to solitary species. (A) Maximum likelihood phylogenetic trees of seven spider species reconstructed by 2824 of core single-copy orthologous genes with 100% ML bootstrap values for all nodes. Divergence time of seven spiders estimated by Timetree (http://www.timetree.org/). The node bars indicate 95% posterior probability intervals. (B) Comparison of number of orthologous gene family in seven spider genomes. Gene family expansion and contraction indicated by plus (+) and minus (-), respectively. (C) Average dN/dS ratios of concatenated all orthologs in seven spider species estimated by branch model in PAML. (D) Violin plot showed the dN/dS ratios of each orthologous genes in seven spider species estimated by branch-site mode in PAML.

We downloaded the curated orthology map of Arachnida from OrthoDB (Kriventseva et al. 2015) which contains 8,805 Orthologous Gene Groups (OGGs) shared by nine Arachnid species with sequenced genomes. Of these 8,805 seed orthologous groups in HaMStR (Ebersberger et al. 2009), we identified the orthologs in each spider species with E-values of less than 10^−20^. We aligned and trimmed the nucleotide sequences of the 8,805 orthologous groups using PRANK (Löytynoja & Goldman 2005) with the parameter “-codon” and MATFF (https://mafft.cbrc.jp/alignment/software/), and trimmed using trimAl (Capella-Gutiérrez et al. 2009) with the parameter “-automated1”.

We identified 1:1, one-to-many, and many-to-many orthologs among all seven spider genomes, and strict 1:1 orthologs (i.e. genes for which only one gene from each species matches the given OrthoDB8 ortholog group). For each 1:1 ortholog pair, we only selected the longest transcript associated with the gene for each pair of species. We ran CAFE (De Bie et al. 2006) to analyze the gene family expansion and contraction based on the above identified 1:1 ortholog pairs, and annotated by gene ontology (GO) with the R software, package topGO (https://bioconductor.org/packages/release/bioc/html/topGO.html). We also identified strict 1:1:1:1:1:1:1 orthologs among all seven spider genomes as single-copy orthologs. We defined orthologs that had more than two homologs as multiple-copy orthologs, and orthologs that had only two homologs as unique paralogs. We also identified the core single-copy orthologs that were shared by all seven spider genomes.

### Genome-wide phylogeny construction and divergence time estimation

We aligned each core single-copy ortholog using MUSCLE v3.8.31 (https://www.ebi.ac.uk/Tools/msa/muscle/) with default parameters and trimmed using trimAl (Capella-Gutiérrez et al. 2009) with parameter “-automated1”. To maximize the information content of the sequences and to minimize the impact of missing data, we filtered the core single-copy orthologs with strict constraints, including length (minimum 200 aa) and sequence alignment (maximum missing data 50% in CDS alignments). Next, we prepared two types of gene datasets after filtering. First, we concatenated all core single-copy genes of each species into one-line sequence as a supergene using a custom python script (genome-scale concatenation-based, supergene). Second, we conducted a genome-scale coalescent-based dataset including 2,824 of core single-copy genes. We used ModelTest2 (Posada & Crandall 1998) to detect the best model for phylogeny construction and then used RAxML 8 (Stamatakis 2014) to build the Maximum Likelihood (ML) tree. We built the ML trees using the two types of datasets (concatenation- and coalescent-based) described above in RAXML, respectively. Finally, we reconstructed the species tree using ASTRAL 4.4.4 (Mirarab et al. 2014).

We generated two datasets from the CDS alignments to estimate the divergence time of each species. One dataset contained the first two partitions, including the first and second codon positions of the sequences. The other dataset contained all three partitions corresponding to all three codon positions in the sequences. We estimated the divergence times under a relaxed clock model using the MCMCTree package in PAML4.7 (Yang 2007), with the “Independent rates model (clock□=□2)” and the “JC69 model”. We used 4,000,000 iterations after a burn-in of 2,000,000 iterations, with other parameters as the default settings of MCMCTree. We ran this analysis twice for each dataset to confirm that the results were consistent between runs. We used the time calibrations from TIMETree (http://www.timetree.org/), a public knowledge-base of divergence times among organisms, demonstrating the high reliability of this molecular clock dating strategy.

### Nucleotide substitution rate estimation

To estimate lineage-specific evolutionary rates for each branch of the phylogeny of the seven spiders, we aligned orthologous protein sequences using MUSCLE v3.8.31 (https://www.ebi.ac.uk/Tools/msa/muscle/), derived nucleotide alignments from protein alignments using PAL2NAL (Suyama et al. 2006), and estimated pairwise dN/dS of nucleotide alignments using the CodeML package in PAML 4.7a (Yang 2007). Specifically, we used the free-ratio model to calculate the ratio of non-synonymous (dN) to synonymous (dS) nucleotide changes separately for each ortholog and a concatenation of all alignments of 2,824 single-copy orthologs from the seven species. Parameters, including dN, dS, dN/dS, N*dN, and S*dS values, were estimated for each branch, and genes were discarded if N*dN or S*dS < 1, or dS >2, following a previous study (Goodman et al. 2009).

### Rapidly evolving gene identification

To identify the rapidly evolving genes (REGs) in social and solitary spiders separately, we built multiple single-copy orthologs datasets for social-solitary comparisons, including one social spider versus the five solitary spiders (fig. 2A) and one solitary spider versus the two solitary spiders (fig. 2B). Each clipped species tree including target spider species were prepared, respectively (fig. 2). We ran the CodeML package to identify REGs of each spider species with corresponding gene sets and appointed species tree separately, with the null model assuming that all branches have been evolving at the same rate and the alternative model allowing the focal foreground branch to evolve under a different evolutionary rate. We used a likelihood ratio test (LRT) in R software (MASS package) with df□=□1 to discriminate between the alternative model and the null model for each single-copy ortholog in the gene set. We only considered the genes as evolving with a significantly faster rate in the foreground branch if the adjusted *P* value□<□0.05 and if the dN/dS in the focal foreground branch was higher than that in the focal background branches (i.e. other six spiders). Finally, we identified the core REGs shared by two social spiders or five solitary spiders, and annotated by GO using the R software, package topGO (https://bioconductor.org/packages/release/bioc/html/topGO.html).

**FIG. 2.**
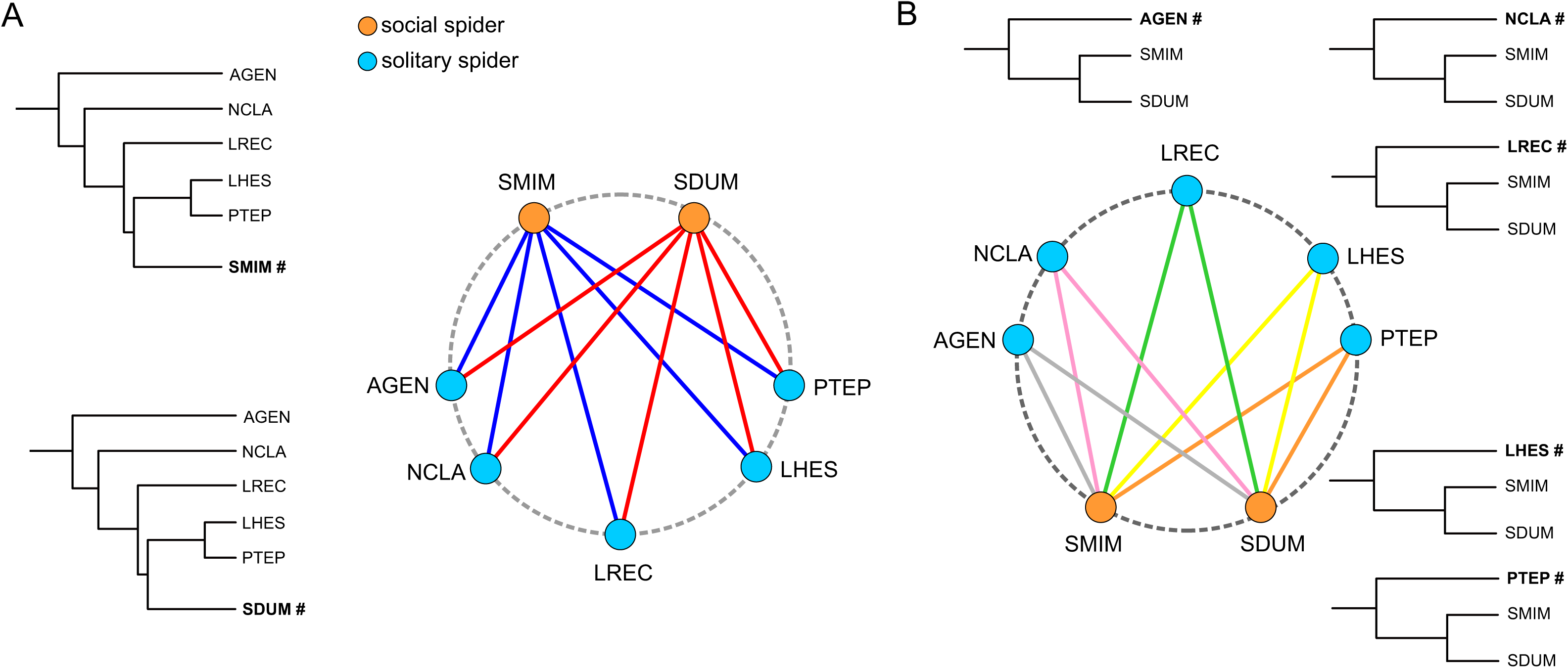
Illustration of comparisons made between social and solitary species to identify rapidly evolving genes (REGs) in social and solitary species. (A) Each of the two social spider species was separately compared to the five solitary spiders and (B) each of the five solitary spider species was separately compared to the two social spiders. Each phylogenetic tree was clipped from the previously reconstructed species tree.

### Expression profiles of rapidly evolving genes in four tissues of the social spider *S. dumicola*

The social spider *Stegodyphus dumicola* is native to central and southern Africa (Johannesen et al. 2007) and has been widely used to study the evolutionary ecology of group living (Pruitt et al. 2019, 2018; Berger-Tal et al. 2014; Grinsted et al. 2013). We collected three colonies of aproximitly 80 individuals each from Kalkrand, Namibia, in February 2017 and housed them in the lab, feeding them with crickets *ad libitum* until RNA extraction. By July 2017, when we extracted the RNA, there were approximately 30-40 individuals in each colony. These individuals were assayed for boldness, by measuring the latency to respond to a simulated predator attack (Keiser et al. 2014; Pruitt & Keiser 2014). Only shy individuals, i.e., those that took 400 sec or longer to recover from the simulated predator attack, were used in this study. All individuals were adult females of the same age, i.e. in their final molt and over a year old. We haphazardly selected 10 shy individuals from each colony for the analysis, resulting in a total of 30 individuals. Individuals were decapitated immediately before tissue extraction. We dissected four tissues: brain, venom gland, legs and abdomen from each individual and immediately stored in RNAlater (Sigma, USA). Samples were stored individually at 4 °C overnight for the RNAlater to penetrate the tissues before being finally stored at −80 °C. Subsequently, samples of each tissue were separately pooled according to colony of origin, resulting in three biological replicate pools (i.e. originating from the three replicate colonies), for each of the four tissues.

The total RNA of each pool was extracted using RNeasy kits (Qiagen, CA, USA), and the quality and quantity of RNAs were detected with Nanodrop 1000 (NanoDrop Technologies, DE, USA) and Agilent Bioanalyzer 2100 (Agilent Technologies, CA, USA). Only RNA with high quality (RNA Integrity Number > 7) were used for cDNA synthesis and amplification. Libraries were prepared with the Nextera XT DNA Library Prep Kit (Illumina, CA, USA) using ∼350-bp inserted fragments for transcriptome sequencing as previously described (Tong, Tian, et al. 2017; Tong, Fei, et al. 2017). Libraries were individually barcoded and run on a single lane of an Illumina NovaSeq (Novogene, CA, USA) yielding 150-bp paired-end reads.

Sequencing reads were checked for quality using Bioconda software, package FastQC. Adapters and reads with a quality score < 20 were trimmed with Trimmomatic (Bolger et al. 2014). We mapped all the clean reads to the *S. dumicola* genome (Liu et al. 2019) using RSEM (Li & Dewey 2011) to obtain expected counts and transcripts per million (TPM). We removed genes with low expression that did not meet one of two criteria: (1) counts per million (TPM) greater than one in at least half the samples, or (2) TPM > 1 in all samples of a given tissue. To classify genes by their tissue specificity, we calculated τ, a commonly used metric of expression specificity (Yanai et al. 2005). τ ranges from 0 to 1, where 0 indicates that genes are ubiquitously expressed and 1 indicates that genes are exclusively expressed in one tissue (Warner et al. 2019). Finally, we compared the expression profiles of REGs across the four tissues, and identified the broadly expressed and tissues-specific REGs.

## Results

### Genome content and evolution

We identified 8,805 Orthologous Gene Groups (OGGs) belonging to Arachnida according to the curated orthology map from OrthoDB (https://www.orthodb.org/). The social spiders S. *mimosarum* (N= 27,135) and *S. dumicola* (N = 37,601) had the most protein coding genes and the largest number of OGGs (N = 7,780, 8,516) relative to the other solitary spider species (Table 1). In addition, even though the two social spiders had fewer orthologs than the solitary species, they had more single-copy orthologs but fewer multiple-copy orthologs (Table 1). The two solitary spider species *P. tepidariorum* and *N. clavipes* had the most orthologs (N = 12,763, 12,784) and multiple-copy orthologs (N = 2,654, 2,579).

**Table 1.** Summary of the gene content of seven spider genomes.

| Species | Sociality | Genes | Orthologous gene groups | Orthologs | Average ortholog number per group | Single-copy orthologs | Multiple-copy orthologs |
| --- | --- | --- | --- | --- | --- | --- | --- |
| *A. geniculata* | Solitary | 23,277 | 5,933 | 9,940 | 1.6754 | 3,682 | 2,251 |
| *L. hesperus* | Solitary | 17,364 | 5,504 | 6,429 | 1.1681 | 4,994 | 508 |
| *L. reclusa* | Solitary | 20,617 | 5,064 | 6,755 | 1.3340 | 4,387 | 6,77 |
| *N. clavipes* | Solitary | 20,241 | 5,895 | 12,763 | 2.1651 | 3,241 | 2,654 |
| *P. tepidariorum* | Solitary | 27,515 | 7,002 | 12,784 | 1.8258 | 4,423 | 2,579 |
| *S. mimosarum* | Social | 27,135 | 7,590 | 7,780 | 1.0250 | 7,540 | 50 |
| *S. dumicola* | Social | 37,601 | 7,118 | 8,516 | 1.1964 | 6,366 | 752 |

In the two social spider genomes, we detected gene family expansions for 214 families in *S. dumicola* and 686 families in *S. mimosarum*, while the five solitary spider species tended to have many fewer gene family expansions and more gene family contractions relative to their most recent common ancestor (fig. 1B). In the *S. dumicola* genome, these expanded gene families were significantly enriched for 128 related gene ontology (GO) categories, mainly consisting of two large groups (supplementary table S2). The first group was related to transport function, such as transmembrane transport (GO:0055085, *P*□=□0.00725), water transport (GO:0006833, *P*□=□0.00456), calcium ion transmembrane transport (GO:0070588, *P* = 0.0072). The second largest group was associated with metabolic processes, including carbohydrate metabolic process (GO:0005975, *P* = 0.00456) and UDP-glucose metabolic process (GO:0006011, *P* = 0.0000088). We also found similar enrichment for gene families expanded for the second social species, *S. mimosarum* (supplementary table S2).

### Genome-wide phylogeny

We identified 2,824 core single-copy orthologs that were shared by the seven spider species. We constructed four maximum likelihood (ML) phylogenetic trees of the seven spiders based on the concatenated (supergene) and coalesced single-copy orthologs. Each phylogenetic reconstruction method (coalescent-based, concatenation-based) and sequence type (nucleotide or amino acid) generated the same topology with 100% ML bootstrap values for all nodes (fig. 1A, supplementary fig. S1). This strongly supported phylogeny also had similar topology to recent phylogenetic studies on *Stegodyphus* species using multiple nuDNA loci (Settepani et al. 2016; Garrison et al. 2016). In addition, we estimated the divergence times for all nodes on the phylogenetic tree (fig. 1A). The two social *Stegodyphus* spiders were estimated to have diverged approximately 14.5921 million years ago (Mya) with confidence intervals 13.5859 to 16.5332 Mya, and diverged from the remaining solitary spider species about 59 Mya (fig. 1A).

### Genome-wide pattern of nucleotide substitution rate

We identified the nucleotide substitution rates in each spider species based on the 2,824 core single-copy genes using PAML (Yang 2007). We found that the two social spiders *S. dumicola* (dN/dS = 0.138194) and *S. mimosarum* (dN/dS = 0.148917) had elevated terminal genome-wide dN/dS compared to their most recent ancestral branch (dN/dS = 0.031015) (supplementary fig. S2). In contrast, the five solitary spider species did not show elevated terminal genome-wide dN/dS relative to their most recent ancestral branch (supplementary fig. S2). Furthermore, the two social *Stegodyphus* spider species had elevated dN/dS compared to the five solitary spider species based on both concatenation-based (fig. 1C) and coalescent-based gene sets (fig. 1D).

### Rapidly evolving gene repertoire

We identified 89 REGs (*P* < 0.05) that had elevated rates of molecular evolution (dN/dS) in both social spider species relative to the five solitary spider species (fig. 3A). Specifically, we identified a total of 158 REGs in the *S. mimosarum* genome, 155 REGs in *S. dumicola* genome, and 89 of these overlapped (supplementary table S3). Significantly enriched GO functional categories for these core REGs in social spiders mainly included three groups: transport, behavior and immune response processes (fig. 3B). REGs associated with behavioral processes included the steroid receptor seven-up (SVP) involved in social behavior (GO:0035176) and aggressive behavior (GO:0002118), sodium channel protein (SCN) involved in mechanosensory behavior (GO:0007638), and oxidation resistance protein 1 (OXR1) related to adult walking behavior (GO:0007628). Transport-related REGs in the two social spiders included genes functioning in metal ion binding, calcium ion binding and sodium ion transmembrane transport, such as Secretory carrier-associated membrane protein 1 (SCAMP1) and RING finger protein 170 (RNF170) (supplementary table S4). REGs associated with immune response function included toll (TL), CCAAT Enhancer Binding Protein Gamma (CEBPG) and Transcription factor E3 (TFE3).

**FIG. 3.**
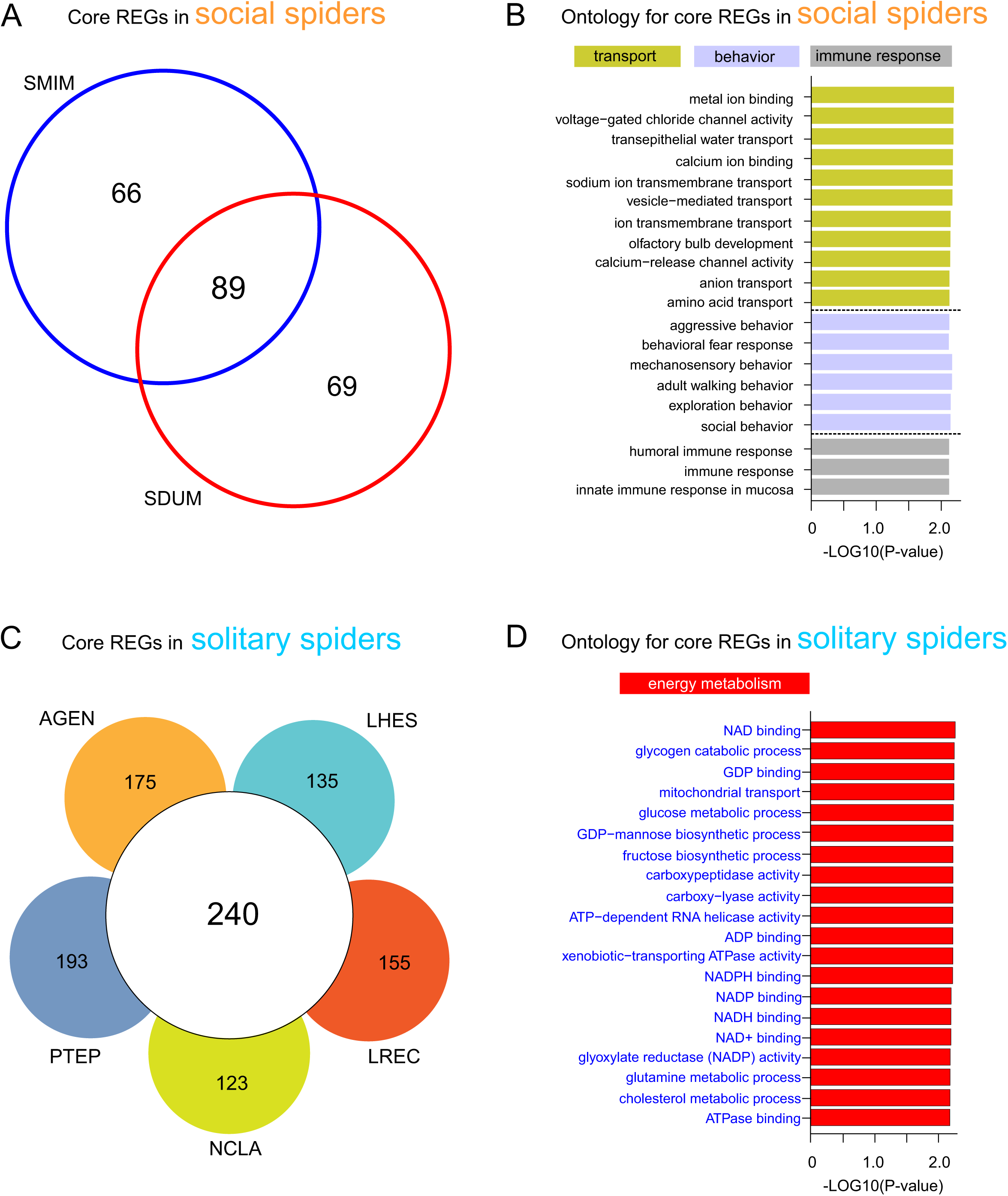
Identification and annotation of shared REGs in social and solitary spider species. (A) Venn diagram showing the number of species-specific and shared REGs in the two social spiders. (B) Bar plot depicting the top 20 gene ontologies significantly enriched in REGs shared between the two social spiders. Ontology for these shared REGs in social spiders was associated with transport, behavior and immune response functions. (C) Venn diagram showing the number of species-specific and shared REGs in the five solitary spider species. (D) Bar plot depicting the top 20 gene ontologies significantly enriched in REGs shared between the five solitary spiders. Ontology for these shared REGs in solitary spiders mainly involved energy metabolism processes.

We identified a total of 240 core REGs shared by the five solitary spider species (fig. 3C, supplementary table S5). More specifically, we identified the largest number of REGs (N = 433) in the *P. tepidariorum* genome and the smallest number of REGs (N = 323) in *N. clavipes* genome. Interestingly, we found that the significantly enriched GO categories of core REGs in the solitary spiders were mainly involved in energy metabolism processes (fig. 3D, supplementary table S6), such as Fatty acid synthase (FAS), Mitochondrial Processing Peptidase Beta Subunit (MPPCB) and Acyl carrier protein (ACP) involved into glucose metabolic process (supplementary table S6).

### Expression pattern of rapidly evolving genes in the social spider *S. dumicola*

Transcriptome sequencing of four tissues (brain, venom gland, legs and abdomen) from the social spider *S. dumicola* generated approximately 86.5 Gb raw reads. After trimming, we obtained nearly 28.6 million PE 150-bp clean reads (supplementary table S7). We focused on the expression patterns between the four tissues of the shared 89 REGs identified in *S. dumicola* and *S. mimosarum* compared to the five solitary spiders (supplementary table S8). Most REGs (N = 81) were expressed in the brain, while fewer REGs (N = 57) were expressed in the venom gland (fig. 4A). We found 53 REGs (59.55%) were broadly expressed in all tissues (fig. 4A). Only a few REGs showed a tissue-specific expressed pattern, including 12 brain-specific genes, three abdomen-specific genes, two leg-specific genes, and zero venom gland-specific genes. Among the broadly-expressed REGs, many of them had higher expression levels in the brain compared to the other three tissues (fig. 4B), such as Glycoprotein 3-alpha-L-fucosyltransferase A (FucTA), AP-2 complex subunit sigma (AP2S1) and AP-1 complex subunit beta-1 (AP1B1). GO enrichment analysis showed that this set of broadly-expressed REGs was enriched in transport processes (fig. 4C, supplementary table S9), such as protein transport (GO: 0015031, *P* = 0.00283), ion transmembrane transport (GO: 0034220, *P* = 0.00323) and ion channel activity (GO: 0005216, *P* = 0.00983). Among the tissue-specific REGs, we found brain-specific REGs had higher expression levels than abdomen-specific REGs (fig. 4D). Notably, significantly enriched GO functional categories of brain-specific REGs included behavioral processes (fig. 4E, supplementary table S10), such as aggressive behavior (GO: 0002121, *P* = 0.00137) and social behavior (GO: 0035176, *P* = 0.00263).

**FIG. 4.**
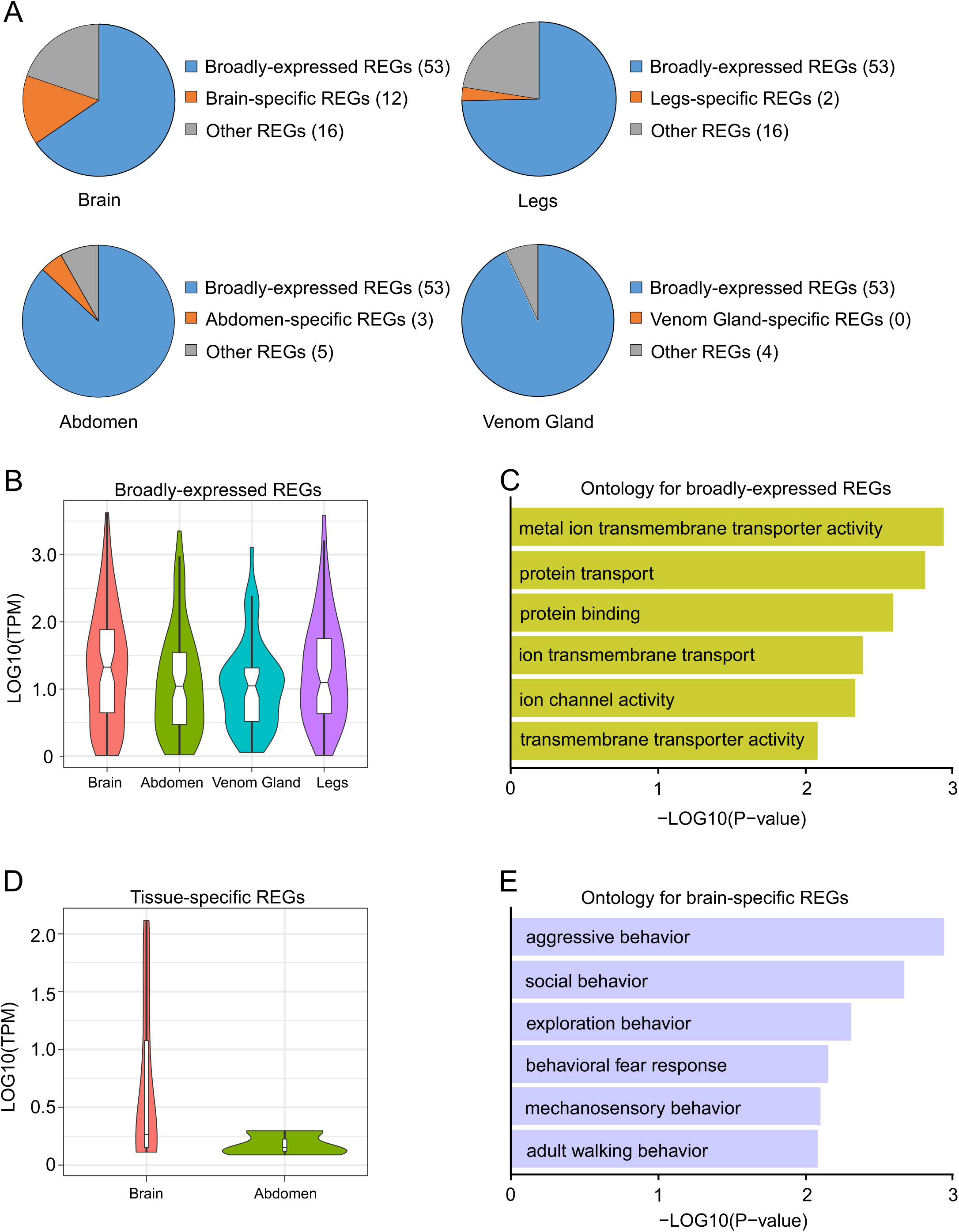
Expression patterns of REGs in *Stegodyphus dumicola*. (A) Distribution of broadly-expressed, tissue-specific and other REGs in each tissue, including brain, legs, abdomen and venom gland. (B) Violin plot depicting the distribution of Log10(TPM) values of broadly-expressed genes labeled by REGs in four tissues. (C) Bar plot depicting the top six gene ontology for broadly-expressed REGs. Ontology for broadly-expressed REGs involved in transport processes (signal transduction). (D) Violin plot depicting the expression ranges of brain- and abdomen-specific genes labeled by REGs. (E) Bar plot depicting the top six gene ontology for broadly-expressed REGs. Ontology for brain-specific REGs involved into social behavior processes.

## Discussion

Building on previous comparative genomic research into the genomic underpinnings of social life in social insects (Simola et al. 2013; Kapheim et al. 2015), we compared the genomes of two social and five solitary spider species. We identified putative genomic signatures of spider social evolution, including expansions of gene families associated with transport and metabolism, and genome-wide elevated rates of molecular evolution in the two social species relative to the five solitary species. Furthermore, we found specific changes in the rate of molecular evolution between the social and solitary species for genes with transport, behavior, immune, and metabolism function. Finally, we found that most rapidly evolving genes in the social spider *S. dumicola* were broadly expressed across four tissues and enriched for transport (e.g., signal transduction) function, but 12 of the rapidly evolving genes in *S. dumicola* showed brain-specific expression and were enriched for genes annotated for behavioral processes.

### Extensive gene family evolution in *Stegodyphus* social spiders targets transport and metabolism functions

Gene families are sets of paralogs that often display similar gene functions. Expansions (gene gain) or contractions (gene lose) in gene families may correspond to adaptive events coupled to life-history transitions (Ranson et al. 2002). We found that the social spiders *S. mimosarum* and *S. dumicola* had large expansions of gene families compared with the five solitary spider species with available genomes (fig. 1A). These extensive gene family expansions of the two social spider species in particular involved gene families associated with metabolism (e.g. carbohydrate metabolic process) and transport (e.g. transmembrane transport) functions. This finding suggests that large-scale transport and metabolism function associated gene duplication (or perhaps whole genome duplication) could be associated with social evolution in spiders.

Previous studies in social insects (ants and bees) identified expansions of gene families associated with metabolism and chemical communication functions (Simola et al. 2013; McKenzie et al. 2016; Wurm et al. 2011; Kapheim et al. 2015). Similarly, previous comparative studies across animals have emphasized genes associated with metabolism as being important in the response to social challenges (Rittschof et al. 2014; Saul et al. 2019), suggesting that changes in genes with metabolic functions might often be involved in social evolution across animals. Indeed, previous research indicates that social spiders may have lower metabolic rates than their solitary relatives (Zimmerman 2007), further implicating changes in metabolic-related genes to the evolution of social life in spiders. Unlike for social insects, we did not detect expansions of chemosensory gene families in the two social spiders. While previous studies have identified large changes in chemosensory gene families across chelicerates (Vizueta et al. 2018, 2017), including spiders, pheromonal communication may be relatively less important for social life in social spiders than in social insects (Zhou et al. 2015; Leonhardt et al. 2016; Vander Meer et al. 1998).

### Social spider genomes exhibit the distinct genome-wide signature of accelerated molecular evolution

We found an elevated genome-wide rate of molecular evolution (dN/dS) in the two social spider species relative to the five solitary species. Our results are consistent with a previous spider study that compared dN/dS ratios for 13 randomly chosen nuclear loci for three social and seven subsocial *Stegodyphus* species (Settepani et al. 2016). Similarly, four eusocial insects were also found to have higher rates of molecular evolution (dN/dS) compared to solitary insects (Romiguier et al. 2014). Studies very often interpret high dN/dS as a putative sign of positive selection (Privman et al. 2018). However, relaxed purifying selection instead of elevated positive selection can also lead to higher average gene-specific or genome-wide dN/dS. Relaxed purifying selection is expected to be especially common in species with low effective population size (N_e_), and social species such as social spiders are expected to experience low N_e_ as a result of reproductive skew, female-biased sex ratios, and inbreeding (Romiguier et al. 2014; Galtier et al. 2018; Settepani et al. 2016). Indeed, previous studies in social spiders (Settepani et al. 2016; Bechsgaard et al. 2019; Settepani et al. 2017, 2014) and also social insects have found evidence for low N_e_ and genome-wide relaxed purifying selection when compared to solitary species (Galtier et al. 2018; Kapheim et al. 2015; Romiguier et al. 2014). Future studies using both polymorphism data and divergence data will be necessary to further tease apart the contribution of elevated positive selection and relaxed purifying selection (Yang & Bielawski 2000; Nielsen 2005) to spider genome evolution.

### Divergent signatures of rapidly evolving genes between social and solitary spiders

Genes that were rapidly evolving (rapidly evolving genes, REGs; i.e. genes with elevated dN/dS) in the two social spider species relative to the five solitary species were significantly enriched in the functions categories associated with transport, social behavior, and immune response. These include genes such as sodium channel protein (SCN), which has previously been implicated in neuronal function and behavior (Ren 2011; Miller 2013). Genes involved in immune response have also been emphasized in social insect studies and may be generally important for the evolution of group-living (Sadd et al. 2005; Vojvodic et al. 2015; Harpur & Zayed 2013). In addition, a set of genes associated with social behavior have been highlighted in social insects and vertebrates (Robinson et al. 2008).

Genes that were rapidly evolving in each of the five solitary spider species relative to the two social species were enriched for energy metabolism function, which together with our gene family expansion results described above, indicate that changes in metabolism may be involved in the evolution of spider sociality. Previous comparative studies have also identified elevated rates of molecular evolution for metabolism-associated genes in eusocial relative to solitary bees (Woodard et al. 2011), and in ants (Roux et al. 2014), although these studies find evidence for higher dN/dS for metabolic genes in more highly social species, while we found the opposite pattern. Altogether, these previous results together with our results suggest that the evolution of social life may often involve changes in genes associated with metabolic function.

### Social spider brain-specific REGs exhibit distinct signature of social behavior bias

To gain insight into the expression pattern of REGs shared in the two social spiders, we characterized the expression profiles of four tissue types (brain, venom gland, abdomen, legs) in *S. dumicola.* Most REGs were broadly expressed across all four tissues, suggesting that genes experiencing rapid molecular evolution in the two social spiders may have general functions. These broadly expressed REGs were enriched for transport-associated functions (e.g., signal transduction) and tended to be more highly expressed in the brain. Most REGs that showed tissue specific expression were found in the brain. Interestingly, these brain-specific REGs were enriched for behavior annotations (e.g., aggressive behavior, social behavior, mechanosensory behavior) (fig. 4, supplementary table S10), and could be candidates for genes influencing social behavior that were involved in the evolution of spider sociality.

### Conclusions

The differences we observed between the genomes of the two *Stegodyphus* social spiders and five solitary spiders are putatively causes or consequences of spider sociality, including both adaptive and non-adaptive evolutionary processes. Alternatively, these differences between the genomes of the two *Stegodyphus* social spiders and the genomes of the other five solitary spiders could also be associated with lineage-specific adaptation or non-adaptive evolutionary processes. In order to further clarify the genomic underpinnings of social life in the spiders, and to disentangle adaptive and non-adaptive genomic signatures associated with spider social evolution, future studies will need to combine more genomic and transcriptomic data from species representing more independent origins of sociality using formal phylogenetic comparative methods (Linksvayer & Johnson 2019; Cornwell & Nakagawa 2017; Garamszegi 2014). Such future studies can confirm whether the putative genomic signatures of spider sociality we found (expansion and rapid evolution of genes with transport, metabolic and behavioral functions, and overall elevated rate of molecular evolution) are consistently found across independent origins of sociality, and can provide further evidence concerning the putative candidate genes we identified.

## Date Archiving

The sequencing reads have been deposited at NCBI SRA under the NCBI BioProject accession PRJNA575239.

## Acknowledgments

We would like to thank Yiyong Zhao of Fudan University for generously providing scripts for phylogenomic analysis. We thank Weitao Chen of Chinese Academy of Fishery Sciences for assistance in molecular clock analysis. We thank the Linksvayer lab members for helpful comments and discussion on an early version of the manuscript. This work was funded by the National Institutes of Health grant GM115509.

